# *Bicaudal C* is required for the function of the follicular epithelium during oogenesis in *Rhodnius prolixus*

**DOI:** 10.1101/2020.02.14.949222

**Authors:** Agustina Pascual, Emiliano S. Vilardo, Catalina Taibo, Julia Sabio y García, Rolando Rivera Pomar

## Abstract

The morphology and physiology of the oogenesis have been well studied in the vector of Chagas disease *Rhodnius prolixus*. However, the molecular interactions that regulate the process of egg formation, key for the reproductive cycle of the vector, is still largely unknown. In order to understand the molecular and cellular basis of the oogenesis we examined the function of the gene *Bicaudal C* (*BicC*) during oogenesis and early development of *R. prolixus*. We show that *R. prolixus BicC* (*Rp-BicC*) gene is expressed in the germarium, with cytoplasmic distribution, as well as in the follicular epithelium of the developing oocytes. RNAi silencing of *Rp-BicC* resulted in sterile females that lay few, small, non-viable eggs. The ovaries are reduced in size and show a disarray of the follicular epithelium. This indicates that *Rp-BicC* has a central role in the regulation of oogenesis. Although the follicular cells are able to form the chorion, the uptake of vitelline by the oocytes is compromised. We show evidence that the polarity of the follicular epithelium and the endocytic pathway, which is crucial for the proper yolk deposition, are affected. This study provides insights into the molecular mechanisms underlying oocyte development and show that *Rp-BicC* is important for de developenta of the egg and, therefore, a key player in the reproduction of this Chagas disease vector.

**Author summary:** The oogenesis is the process of egg formation. It is essential to guarantee transgenerational inheritance. It implies the differentiation of the gamete (oocyte) from a niche of stem cells in the germ line, the accumulation of yolk, and the formation of the chorion. These events are entangled in a regulated manner by the concerted communication between the different cell types that form the ovary. It is regulated by endogenous gene networks and linked to the physiological state of the insect by hormonal clues. This timely orchestrated process represents the interaction of gene networks. The genetic regulation behind the oogenesis is largely unknown in *Rhodnius prolixus*. Here we identified a gene required for egg formation that interferes the uptake of the yolk by affecting the functional integrity of the follicular epithelium. Our results are of interest for a better understanding of a complex process essential for the survival of vector populations and provide knowledge to envisage and design new strategies for vector control.

## Introduction

*Rhodnius prolixus*, is a hematophagous insect and, like other triatomines, is the vector of *Trypanosoma cruzi*, the agent of Chagas disease [1,2]. Chagas disease is a life-threatening disease affecting millions of people worldwide [3]. As vaccine are unavailable and disease treatment is unsafe, vector control is still the most useful method to control the illness. In this context, as oogenesis is crucial for embryo viability and population dynamics, molecular investigation on this process could represent an interesting target to develop novel strategies for insect population control [4,5].

In addition to the sanitary relevance, *R. prolixus* has been a classical model of physiology since the pioneer studies of Sir Vincent Wigglesworth [6–9] and, to some extent, an emerging model for developmental biology [10–12]. The genome of *R. prolixus* has been sequenced [13] and several tissue-specific transcriptomes have been reported [14,15], providing a solid foundation for gene identification. The development of molecular tools such as parental RNA interference (RNAi) [11] set the ground for functional analysis.

The formation of the egg, namely oogenesis, is a period of rapid cellular growth and differentiation which is triggered by feeding. The oogenesis implies the differentiation of the oocyte from a niche of stem cells in the germ line, the accumulation of yolk, formation of the chorion, and the establishment of the future embryo axes. These events are entangled in a regulated manner by means of the communication between the different cell types that compose the ovary, hormonal signaling, that modulate the action of gene networks [16]. Insect ovaries are classified into three distinct types: panoistic, polytrophic and telotrophic, based on the morphology of germ cells in the mature ovary [17–21]. The morphology and architecture of the ovaries have been studied in a variety of insects [4,21–27], but the complete regulatory profiles of gene expression have only been determined in *Drosophila melanogaster* [28–33].

The adult females of *R. prolixus* have two ovaries, each one made up of seven ovarioles [34]. The ovaries are telotrophic, in which nurse cells are confined to a distal chamber referred to as the trophic chamber or tropharium, separated from the vitellarium, structure in which oocytes go through the different stages of oogenesis, previtelogenesis, vitelogenesis and choriogenesis, accompanied by the follicular epithelium [26,35]. The trophic chamber produces maternal RNAs and nutrients, which are transported to the developing oocyte through tubular bridges-the trophic cords, in a directional transport mediated by a network of microtubules [36,37]. The accompanying follicle cells shows dramatic morphological and physiological changes during the different stages of oogenesis, but they always keep an organized pattern [26]. Together with the fat body, the ovaries are responsible for the synthesis of vitellogenin (Vg), precursor of vitellin (Vn), the main component of egg yolk [38,39]. Later on, follicle cells produce the outermost layer of the egg, the chorion, which protects egg from dehydration and regulates oxygen intake and fertilization [40]. Despite the detailed studies [41], we still lack information about the gene networks involved in *R. prolixus* oogenesis.

Many orthologues of the genes involved in *D. melanogaster* oogenesis has been identified in *R. prolixus* [13,14,42]. One of this was *Bicaudal C (BicC). BicC* was originally identified in a *D. melanogaster* maternal mutagenesis screen [43–45]. Females heterozygous for *BicC* mutations produce embryos of several different phenotypic classes, including bicaudal embryos that consist only of a mirror-image duplication of 2–4 posterior segments. Homozygous *BicC* females are sterile because the centripetal follicle cells fail to migrate over the anterior surface of the oocyte at stage 10 during *D. melanogaster* oogenesis [44–46]. *BicC* encodes a protein with hnRNP K homology (KH) and sterile alpha motif (SAM) domains [46], both RNA-binding motifs [47–49]; that interacts with other proteins related to RNA metabolism and targets mRNAs to form regulatory ribonucleoprotein complexes [50]. Also, it has been reported to be involved in the function of Malpighian tubules in the adults [51].

Here, we report the function of *Bicaudal C (Rp-BicC*) during oogenesis of *R. prolixus*. We identified the expression of *BicC* gene and carried out parental RNAi experiments. Our results show that *Rp-BicC* is required for the proper follicle cell function in early stages of oogenesis, affecting yolk uptake, but not choriogenesis.

## Materials and methods

### Insect husbandry

A colony of *Rhodnius prolixus* was maintained in our insectarium of the Centro de Bioinvestigaciones (CeBio) in plastic jars containing strips of paper at 28°C and 80% relative humidity in controlled environment incubators with a 12h light/dark cycle. In this condition, embryogenesis takes 14 ± 1 days. Insects were regularly fed on chicken, *ad libitum*, which were housed, cared, fed and handled in accordance with resolution 1047/2005 (National Council of Scientific and Technical Research, CONICET) regarding the national reference ethical framework for biomedical research with laboratory, farm, and nature collected animals, which is in accordance with the international standard procedures of the Office for Laboratory Animal Welfare, Department of Health and Human Services, NIH and the recommendations established by the 2010/63/EU Directive of the European Parliament, related to the protection of animals used for scientific purposes. Biosecurity rules fulfill CONICET resolution 1619/2008, which is in accordance with the WHO Biosecurity Handbook (ISBN 92 4 354 6503).

### Identification of the *BicC* transcript and cDNA synthesis

The transcript was identified by local BLASTX search using the *D. melanogaster* orthologue gene as query on a transcriptome from ovary and different early embryonic stages assembled using the annotated genome of *R. prolixus* as reference (VectorBase, RproC3 version; Pascual and Rivera Pomar unpublished data). RNA, was isolated from *R. prolixus* embryos at different pre-gastrulation developmental times using TRIZOL reagent (Invitrogen). cDNA was synthesized using kit SuperScript™ VILO™ MasterMix (Invitrogen) and used as template for PCR. Specific primers for *Rp-BicC* were designed [52,53] to amplify two different regions within the KH domain *Rp-BicC^1^* (237 bp): sense-1 5’-CAAGGCACGTCAACAGCTAA-3’, antisense-1 5’-GGATCGTTAGGAGCGATCAA-3’; and *Rp-BicC^2^* (291 bp): sense-2 5’-CGACTCAAACTTGGTGCAAA-3’, antisense-2 5’-AACTTCGCCAGCGATAGAAA-3’. The reaction conditions were 5 min at 94°C, followed by 35 cycles of 30 seconds (s) at 94°C, 30 s at 60°C and 35 s at 72°C and a final extension of 5 min at 72°C. Amplicons were separated in 1% agarose gels, and sequenced to confirm identity (Macrogen Inc.). In addition, the same primers were designed containing T7 promoter sequence (CGACTCACTATAGGG) at the 5’end for use for *in vitro* transcription of dsRNA or antisense RNA probes.

### Ovary and embryo manipulation

Control and silenced adult females were fed to induce oogenesis and five days later the ovaries were dissected in Phosphate Buffered Saline (PBS 1X). Ovaries were fixed in different ways depending on the subsequent analysis. For confocal microscopy, the fixation was performed on ice in 4% paraformaldehyde (PFA) in PBT (PBS 1X + 0.1% Tween-20) for 30 minutes (min), then washed three times in PBT and stored at 4°C until staining the nuclei with Hoescht (Sigma-Aldrich, USA, 1 μg/ml). Images were acquired with the Zeiss LSM 800 confocal microscope. For light microscopy ovaries were fixed in formaldehyde 4%, washed with Millonig’s buffer, dehydrated in graded series of ethanol (70%, 96%, 100%) and xylene (100%) and embedded in paraffin [54]. 5 μm thick sections were cut in a rotary microtome (Leica) and stained using standard hematoxylin-eosin procedure, mounted and photographed using an A1 ZEISS microscope. For transmission electron microscopy (TEM), the protocol was modified from Huebner and Anderson [55]. Ovaries were fixed in glutaraldehyde 2.5% and post-fixed in 1% osmium tetroxide in Millonig’s buffer at pH 7.4. This was followed by dehydration in a graded series of ethanol (25%, 30%, 50%, 80%, 90%) and acetone (100%), after which the ovaries were infiltrated and embedded in epoxy resin (Durcupan ACM, Fluka AG, Switzerland). Ultrathin sections (~60 nm) were cut with a diamond knife, stained with aqueous uranyl acetate and Reynold’s lead citrate [56], and examined at 80 kV in a MET JEOL 1200 EXII transmission electron microscope.

For the analysis of lipids and membranes distribution, lipophilic styryl dye FM 4-64FX (Thermo Fisher Scientific) was injected in the body cavity of females in a 1:500 dilution (3 μg/μl) in PBS 1X and let to diffuse for 20 min. FM 4-64FX targets plasma membrane and marks exo/endocytosis hot spots in the cells [57]. For the dextran oocyte uptake analysis, 2 μl of Texas Red-conjugated dextran (10.000 MW; Molecular probes, Thermo Fisher Scientific) was injected between abdominal tergites in the hemocoel of females 4 days after blood meal and incubated 24 h. After the corresponding time, ovaries were dissected and fixed as described above for confocal microscopy and images were acquired in a Zeiss LSM 800 confocal microscope.

Eggs collected from individual females were used for scanning electron microscopy (SEM), fixed in glutaraldehyde 2.5%, washed with Millonig’s buffer, dehydrated in a graded series of ethanol (70%, 96%, 100%), mounted with double-sided adhesive carbon tape on metallic stubs, metallized with gold and observed under a SEM Quanta 250 (FEI) operated at 20 kV [58].

### Fluorescent immunohistochemistry

Ovaries were fixed for confocal microscopy, then washed in PBX (0.1% Triton X-100 in PBS), blocked with 5% normal goat serum for 2 h, and incubated overnight at 4°C with 1:200 dilution of rabbit polyclonal anti-vitellin antibody (gamma-globulin fraction) [59,60]. After extensive washing, the ovaries were incubated for 2 h at room temperature with secondary Alexa 568-conjugated antirabbit IgG (1:500 in PBX; Invitrogen, Life Technologies), washed and counterstained with Hoescht (Sigma-Aldrich, USA, 1 μg/ml) before image acquisition in a ZEISS LSM 800 confocal microscope.

### RNA *in situ* hybridization

Digoxigenin-labeled antisense *Rp-BicC* RNA probes were synthesized using the RNA-Dig Labeling kit (Roche). *In situ* hybridization was carried out in 4% PFA fixed ovaries stored in PBT at 4°C. The ovaries were post-fixed in PBT + fixative solution (10% PFA in PBS + EGTA-Na_2_) for 20 min on a rocking platform at room temperature. The ovaries were washed three times with PBT and digested with proteinase K (10mg/ml) for 15 min and post fixed as before, following three PBT washes. A pre-hybridization step was performed for 2 h at 60°C in Hybe (50 % formamide, 5x SSC, 0.2 mg/ml Sonicated salmon testes DNA, 0.1 mg/ml tRNA, 0.05 mg/ml Heparin, 0.1 % Tween-20) before the addition of the probe and further incubated overnight at 60°C. The ovaries were rinsed three times with Hybe-B (50 % formamide, 5x SSC, 0.1 % Tween-20) and then washed in Hybe-C (50 % formamide, 2x SSC, 0.1 % Tween-20) during 2h at 60°C, and further washed three times with PBT. The hybridized samples were blocked with antibody-hybridization solution (0.2% Tween-20, 1 mg/ml Bovine Serum Albumin, 5% Normal Goat Serum) for 3 h at room temperature and then incubated overnight with alkaline phosphatase-conjugated anti-DIG Fab fragments (Roche, 1: 2,000) at 4°C on a shaking platform. The antibody was washed away three times with PBT and one time with alkaline staining buffer (100 mM TRIS, 100 mM NaCl, 0.1% Tween-20). The enzymatic activity revealed with NBT/BCIP (Roche). When staining was evident, the ovaries were washed in PBT three times to stop the reaction, dehydrated in a graded series of ethanol and mounted in glycerol for observation and image acquisition using A1 ZEISS microscope.

### Parental RNA interference

Double-stranded RNA (dsRNA) was produced by simultaneous transcription with T7 RNA polymerase (New England Biolabs) on PCR products containing T7 promoter sequences (CGACTCACTATAGGG) at both ends. Two independent templates, dsRNA^*BicC*1^ and dsRNA^*BicC*2^ were used for independent experiments to evaluate potential off-target effects. dsRNA was quantitated and injected into virgin females, using different concentrations, as described in Lavore et al. [11]. Two days after injection, the females were fed to induce oogenesis and mated with males. After mating, eggs were collected and ovaries fixed as indicated above. A negative control was performed injecting virgin females with dsRNA corresponding to the β-lactamase gene (dsRNA^*β-lac*^) of *Escherichia coli* gene [11].

## Results

### The *Rp-BicC* transcript is expressed in ovaries and early embryos

*Rp-BicC* was identified in transcriptomes derived from ovaries, unfertilized eggs and early embryos of *R. prolixus* (Pascual and Rivera-Pomar, unpublished data) by sequence similarity search against *D. melanogaster* orthologue. The assembled *Rp-BicC* transcript from these RNA-seq data set corresponds to the *ab initio* annotated transcriptional units RPRC0001612 and RPRC001613 within the supercontig KQ034133, indicating that the two different predictions in Vector Base were erroneous, and correspond to an only transcriptional unit (**Fig. 1A**). The transcript (1,986 bp) derives from 14 exons and encodes a predicted polypeptide of 662 amino acids. Multiple alignment of *BicC* orthologous sequences showed that *Rp-BicC* conserve the typical KH and SAM domains as other species (**Fig. 1B** and **Fig. S1**).

**Fig 1.**
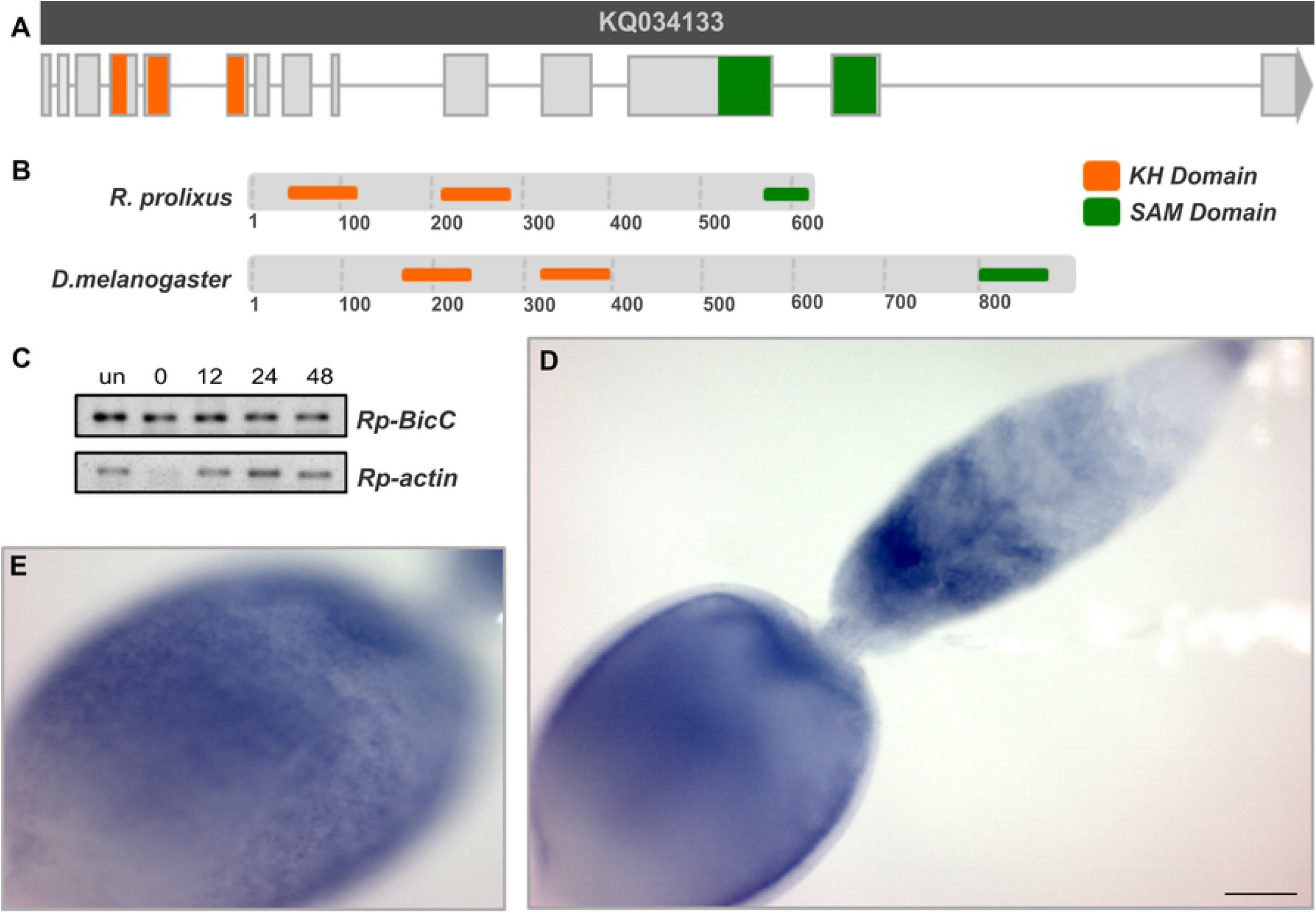
Structure and expression of *R. prolixus Bicaudal C (Rp-BicC*). (**A**) Scheme of the gene structure. Grey bar represents the supercontig that contains the *Rp-BicC* transcriptional unit. Light gray boxes represent exons. (**B**) Diagram of the predicted conserved functional domains of *BicC* in *R. prolixus* and *D. melanogaster* domains. (**C**) Detection of *Rp-BicC* transcript in the ovarioles by *in situ* hybridization. The arrowhead indicates the expression in distal part of the tropharium, the arrow indicates the expression in the vitellogenic oocyte. Scale Bar: 100 μm. (**D**) Different focal plane of the vitellogenic oocyte showed in **C**. Note the expression of *Rp-BicC* in the follicular cells. (**E**) Detection of *BicC* transcript by RT-PCR at different developmental, unfertilized eggs (un), 0, 12, 24, 48 hours post egg-laying (hPL). Upper panel, *Rp-BicC*, lower panel, *Rp-actin*.

Reads from the *Rp-BicC* transcript were identified in all of the transcriptomes corresponding to the different stages come from ovaries, unfertilized eggs and early embryos at 0, 12, 24 and 48 hours post egg-laying (hPL; Pascual and Rivera-Pomar, unpublished data), indicating that the transcript is maternally contributed, although the zygotic expression cannot be ruled out. The expression was assessed by RT-PCR in unfertilized (un), early zygote (0 hPL), blastoderm (12 hPL), gastrulating germ band (24 hPL) and germ band (48 hPL) eggs. *Rp-BicC* mRNA was detected in all stages analyzed (**Fig. 1B**). *In situ* hybridization revealed expression of the *Rp-BicC* transcript in ovaries, showing cytoplasmic distribution in both, the germarium (**Fig. 1D**, arrowhead) and the follicular epithelium of previtellogenic and vitellogenic oocytes (**Fig. 1D, E**).

### *Rp-BicC* is required for proper egg formation

To determine the role of *Rp-BicC*, we injected non-fed virgin females with different concentrations of two independent dsRNA corresponding to different regions of the transcript (dsRNA^*BicC1*^, 237 bp and dsRNA^*BicC2*^, 291 bp). As control, we used dsRNA corresponding to the *β-lactamase* gene of *E. coli* (dsRNA^*β-lac*^). After feeding and mating, dsRNA^*Rp-BicC1*^, dsRNA^*BicC2*^, and dsRNA^*β-lac*^ injected females were evaluated for fertility, egg deposition and morphology, and embryonic and ovary phenotype. The silenced females laid fewer eggs than the control, suggesting that fertility is compromised (**Table S1**). The eggs were let to develop for the expected time of embryogenesis to finish (>14 days), but none of the eggs from interfered females resulted in hatchlings, indicating that the embryogenesis was affected. Dissection of the eggs showed that they lack any distinguishable embryonic structure, suggesting that *BicC* might act at very early stages of development (data not shown).

The eggs laid by the silenced females, as opposed to the control ones, were smaller, with irregular shape and presented white coloration instead of the characteristic pink (**Fig. 2A**), indicating the absence or significant reduction of the *Rhodnius heme-binding protein* (RHBP, one important component of the yolk). The *Rp-BicC* interfered eggs showed an irregular surface. To determine if there is a structural alteration in the chorion morphology we performed scanning electron microscopy. Compared to the regular hexagonal pattern of the chorion observed in the control (**Fig. 2B**), the eggs derived from the silenced females showed defects in the chorion structure, showing an irregular pattern, prominences, and a shrink surface (**Fig. 2C**). The operculum is deformed, although it has a similar size as the control ones. This indicates that the chorion and chorion structures are formed, but the regular patterning is dramatically affected.

**Fig 2.**
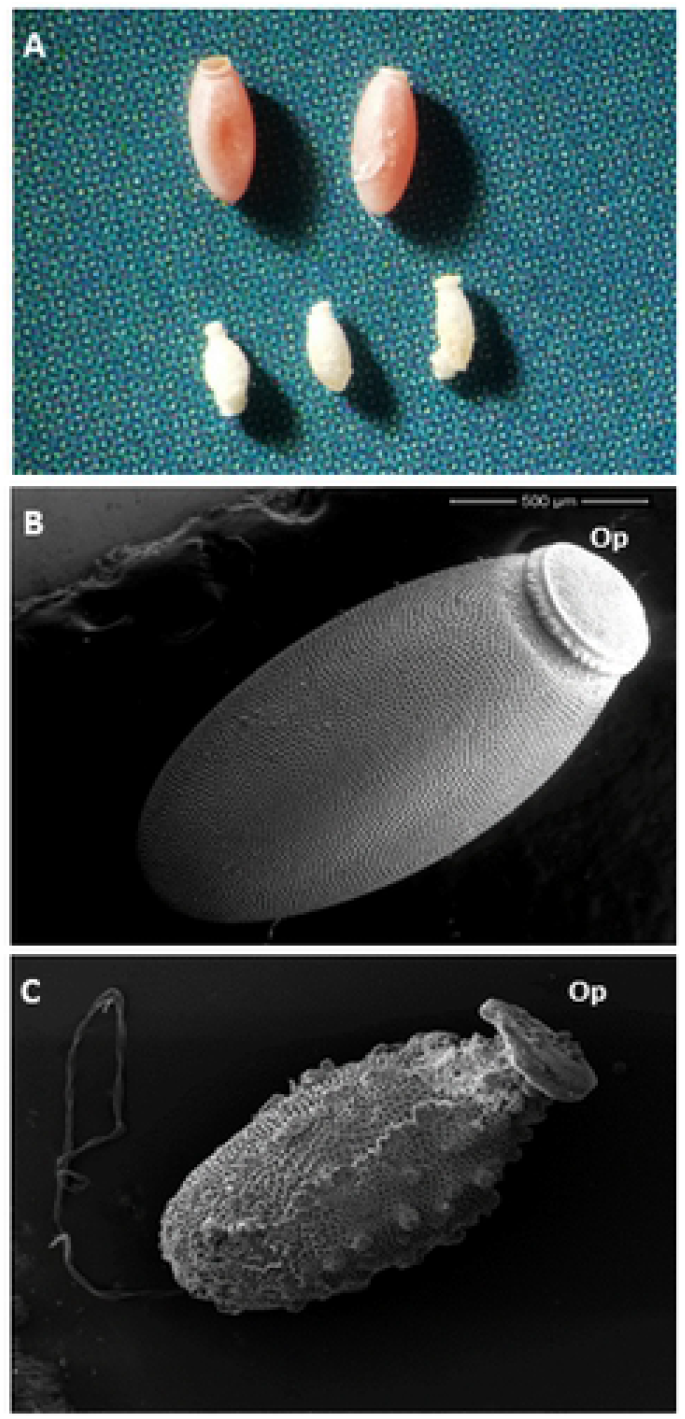
Silencing of *Rp-BicC* results in defective chorion formation. (**A**) Eggs from control (upper row) and silenced (lower row) females observed with a dissecting microscope. Note the smaller size and the lack of the characteristic pigmentation of the eggs from silenced females. (**B**) Scanning Electron Microscopy image of a control egg. (**C**) Scanning Electron Microscopy image of an egg from interfered female. Scale bar: 500 μm. Op, operculum, it corresponds to the anterior pole of the egg.

### *Rp-BicC* is required for the development of follicular epithelium during oogenesis

To further investigate the effect of *Rp-BicC*, we studied the morphology of the ovary. The ovaries of the *Rp-BicC* silenced females have the same number of ovarioles as the control, but they are reduced in size (**Fig. 3A-F**). We analyzed the morphology under DIC optics (**Fig. 3B, E**) and by staining the nuclei to determine cell distribution (**Fig. 3C, F**). Compared to the control (**Fig. 3B**), the follicular epithelium of the *Rp-BicC* silenced females was folded and wrinkled (**Fig. 3C**) and both, previtellogenic and vitellogenic oocytes were smaller (**Fig. 3B-C**). The ovaries of *Rp-BicC* silenced females did not evidence significant morphological differences in the germarium, but displayed the absence of the large nucleoli characteristic of the trophic chamber. From previtellogenic stages on, we observed that the regular organization of the follicular cells is lost (**Fig. 3E, F**). Thin sections of the ovary stained with hematoxylin/eosin showed that, as compared to the control ones (**Fig. 3D**), silenced females displayed oocytes with irregular yolk distribution, accompanied by diminished number of yolk granules and presence of empty spaces in the cytoplasm. The follicular cells appear detached one from each other and the irregular columnar epithelium showed increased intercellular space (**Fig. 3H**). Transmission electron microscopy analysis indicates that, compared to the control (**Fig. 3I**), follicular cells of the *Rp-BicC* silenced females lack their contact with the basal membrane and tunica propria, reduction of the contacts that keep them together in a regular manner, and show vesiculated cytoplasm and less dense nucleoli. (**Fig. 3J**). This results agrees with the observed phenotype of the chorion and indicate that the follicular cells are able to form the chorion, despite the disarray of the cells.

**Fig 3.**
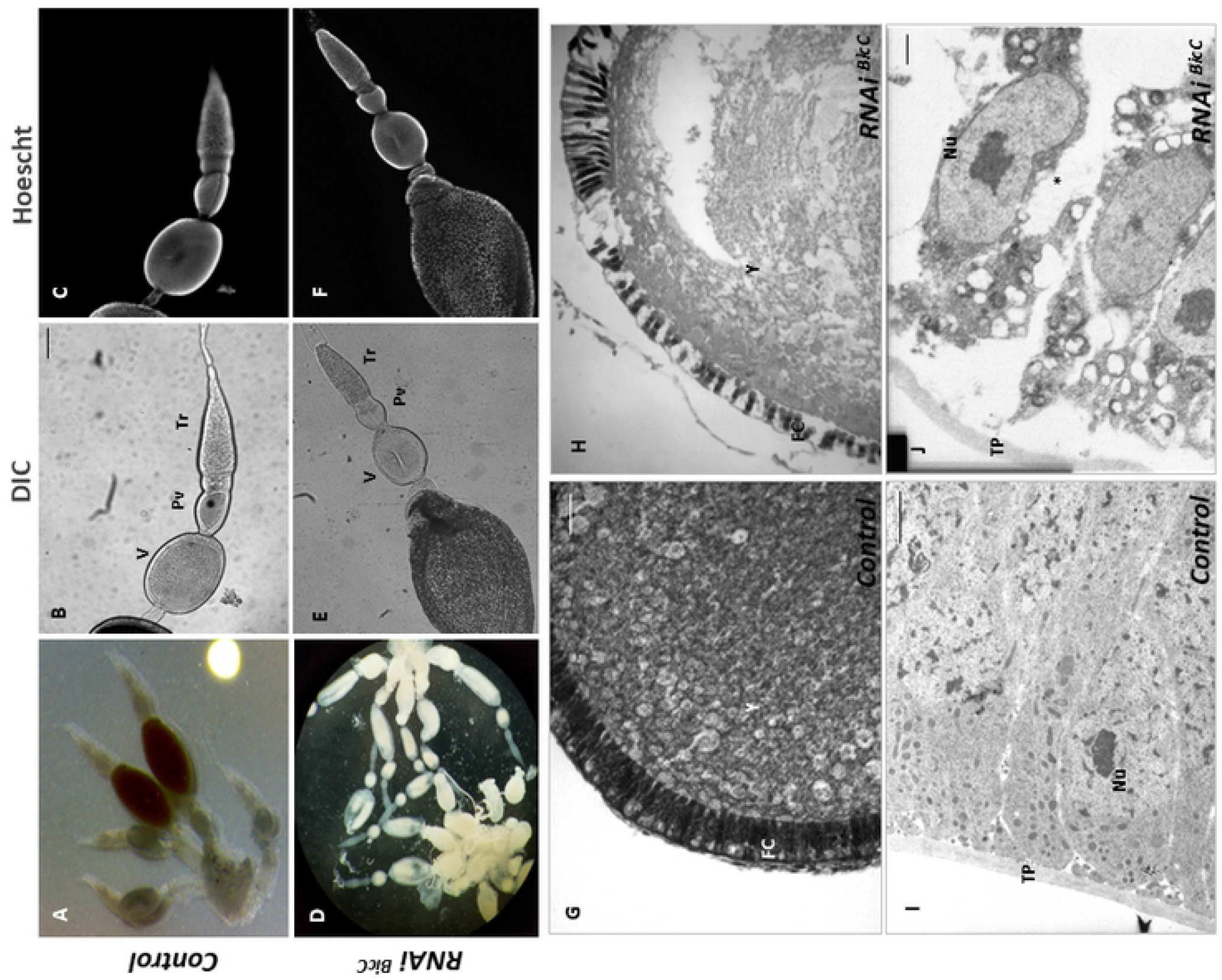
Silencing of *Rp-BicC* affects the ovary morphology. (**A**) Ovary morphology of a control female under dissecting microscope. (**B-C**) Ovariole of a control female showing the tropharium (Tr) and a previtellogenic (Pv) and vitellogenic (V) oocytes by differential interference contrast microscopy (**B**) and nuclei distribution by Hoescht staining (**C**); Scale bar: 200 μm. (**D**) Ovary morphology of a RNAi^*BicC*^ female under dissecting microscope. Note the smaller size of the ovarioles and the lack of pigmentation. (**E-F**) Ovariole of a silenced female showing the tropharium (Tr), a previtellogenic (Pv) and vitellogenic (V) oocytes by differential interference contrast microscopy (**E**) and nuclei distribution by Hoescht staining (**F**); Scale bar: 200 μm. Note that the smaller size of the previtellogenic oocyte and the disarray of nuclear distribution of the follicular cells. (**G**) Histological staining (Haematoxylin-Eosin) of a vitellogenic oocyte from control females; Scale bar: 10 μm. FC, follicular cells. Y, yolk. (**I**). Transmission electron microscopy (TEM) of a previtellogenic oocyte from control females; Scale bar: 2 μm. TP: Tunica propria. Nu: Nucleolus. (**H**) Histological section (Haematoxylin-Eosin staining) of a vitellogenic oocyte from interfered females; Scale bar: 10 μm. Note the space between follicular cells and the inhomogeneous distribution of yolk. (**J**) Transmission electron microscopy (TEM) of a previtellogenic oocyte from silenced females; Scale bar: 2 μm. The asterisk marks the intercellular space Intercellular spaces.

### *Rp-BicC* affect the polarity and vesicle trafficking of follicle cells

In order to address the functional characterization of the morphological changes observed in the follicular epithelium, we analyzed whether the yolk uptake and the polarity of the follicular cells were affected. We used an antibody to localize the presence of vitellin in the developing oocytes. In control females, we observed that vitellogenic oocytes accumulates vitellin in the follicle cells (**Fig. 4A-C**). The ovaries of silenced females, compared to the control, showed a dramatic decrease of the anti-vitellin signal (**Fig. 4D-F**). A closer look showed that vitellin was concentrated in granules in the apical region of the follicle cells (**Fig. 4G**), while in the silenced females very few granules could be accounted and the signal amount was lower (**Fig. 4H**). We hypothesis that a decreased amount of vitellin in the cell, although we can not rule out a dispersed localization in the silenced females. One reason of this might be that the loading of the vitellin by the follicle cells is affected. Therefore, we used a lipophilic styryl dye to mark cell membrane and nascent endosomes that spread into the cytoplasm. In the ovaries of control females, we observed defined fluorescent signal in the apical and basal poles of the follicle cells indicating *bona fide* regions of endo and exocytosis (**Fig. 4I-K**). In the ovaries of *Rp-BicC* silenced females, the fluorescence could only be detected in the membrane in apical pole (**Fig. 4L-M**). This suggests that the polarity of the follicular epithelium is compromised and, therefore, it might affect the interaction of the follicle cells with the developing oocyte. As we have shown before, the accumulation of yolk drops in the ovaries of *Rp-BicC* silenced females, thus we conclude that the lack of a fully functional endo/exocytic pathway might affects the transport of vitellin to the oocyte. We hypothesized that if the endo/exocytic pathway is affected, there should be a general defect in the transport of molecules from the haemolymph to the oocyte through the follicle cells. To test this, we injected fluorescent dextran (MW 10 kDa) in the abdominal cavity of both, control and silenced females, and analyzed the differences in the uploading of the dextran. In control females, the vitellogenic oocytes accumulates fluorescent dextran (**Fig. 5A**; the general morphology of the ovariole is shown by Hoescht staining in **Fig. 5B**), while the vitellogenic oocytes of the *Rp-BicC* silenced females shows a dramatic reduction of fluorescence (**Fig. 5C**; morphology in **Fig. 5D**). Taken together our results supports the notion that the lack of *Rp-BicC* affects the transport through the follicular epithelium in the ovary.

**Fig 4.**
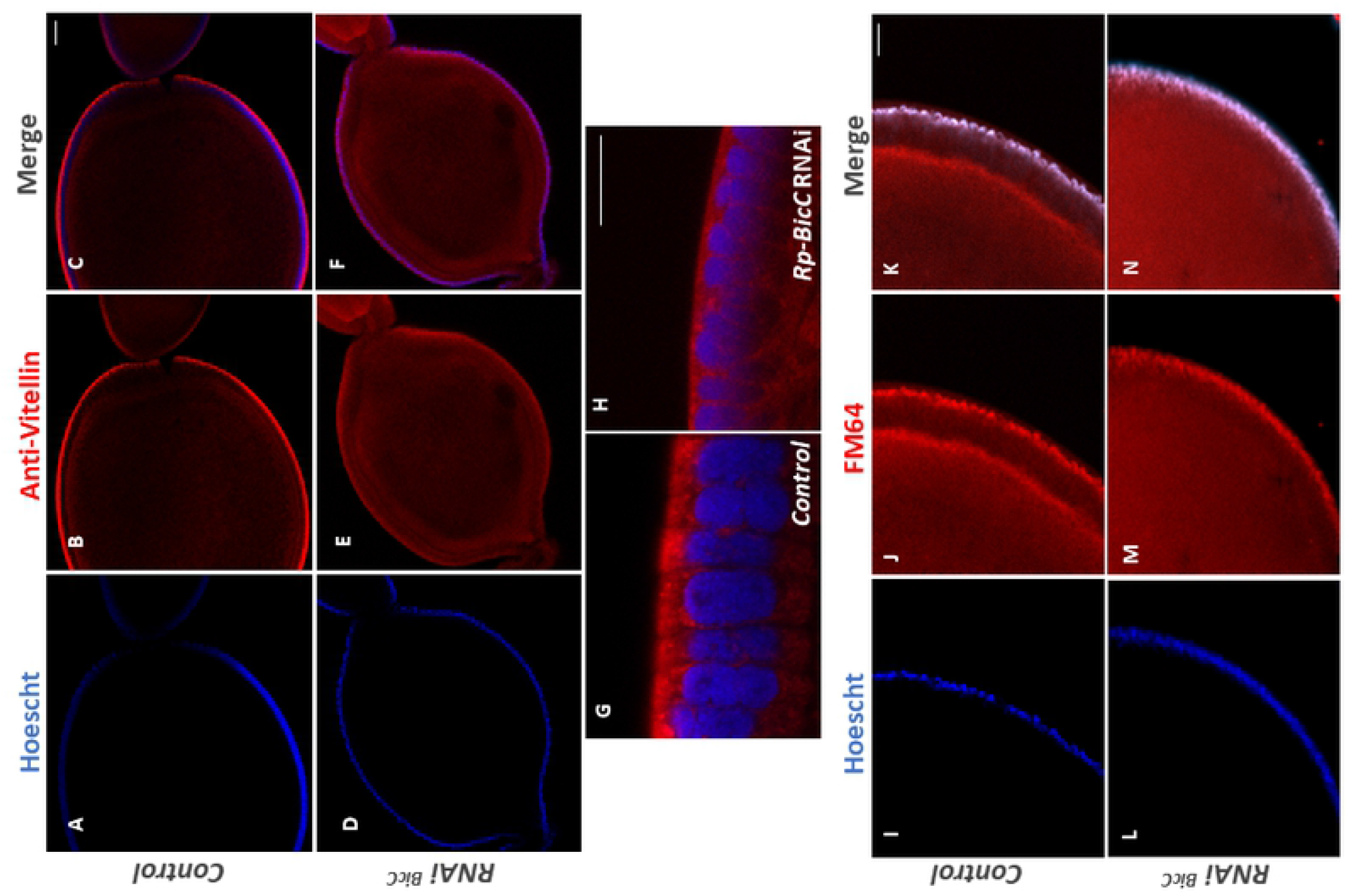
Silencing of *Rp-BicC* alter vitellin and lipid distribution in vitellogenic oocytes. (**A-C**) Immunostaining using anti-vitellin antibody to determine distribution of vitellin (visualized in red) in the oocytes of control females. (**D-F**) Immunostaining using anti-vitellin antibody to determine distribution of vitellin (visualized in red) in the oocytes of RNAi^*BicC*^ females. All samples were counterstained with Hoescht (blue). Scale bar: 50 μm. (**G**) Confocal optical section of anti-vitellin immunostained follicular cells from control females. (**H**) Confocal optical section of anti-vitellin immunostained follicular cells from silenced (RNAi^*BicC*^) females. Scale bar: 10 μm. (**I-K**) Distribution of the FM 4-64FX probe in oocytes after injection of control females. (**L-N**) Distribution of the FM4-64FX probe in oocytes after injection of silenced (RNAi^*BicC*^) females. Hoescht (**I, L**, in blue) visualizes DNA; FM 4-64FX (**J, M**, in red) stain membranes and endocytic vesicles. Scale bar: 50 μm.

**Fig 5.**
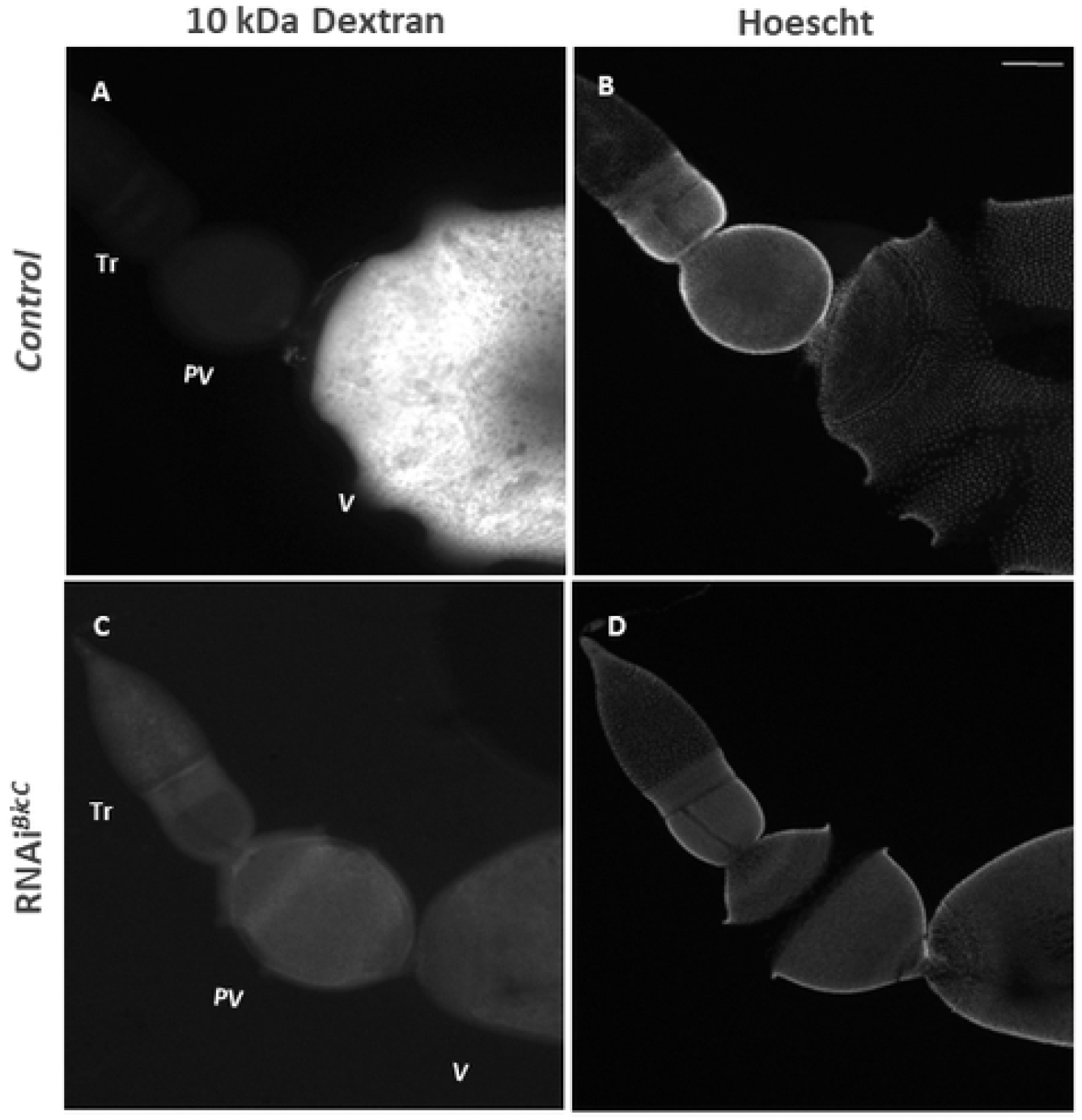
*In vivo* dextran uptake in oocytes is prevented in RNAi^*BicC*^ females. (**A**) Distribution of Texas Red-labeled dextran in vitellogenic oocytes from control females. Tropharium (Tr), Previtellogenic oocyte (Pv), Vitellogenic oocyte (V). Exposure time: 50 msec. (**B**) Counterstaining of control with Hoescht. (**C**) Distribution of Texas Red-labeled dextran in vitellogenic oocytes from silenced (RNAi^*BicC*^) females. Exposure time: 100 msec. (**D**) Hoescht counterstaining of **C**. Due to the large size of the ovarioles of *R. prolixus*, some of them might be squashed by the coverslip after mounting resulting in breaking of the follicular epithelia which is not dependent on the interference experiment. Scale bar: 50 μm.

## Discussion

*R. prolixus* has been a model system for many essential issues in biology, but the understanding of the molecular mechanisms had to wait until the sequencing of the genome [13]. The physiology of *R. prolixus* has been studied since the pioneering work of Vincent Wigglesworth [61,62]. The oogenesis of *R. prolixus* is one of the best studied among insects, from the morphological work of Huebner [22,24–26,34,63] to the biochemistry studies of Masuda [64–70]. Although studies on the cellular biology of oogenesis have recently emerged [71,72], we still lack enough information on genetic to understand the molecular basis of egg formation and patterning in *R. prolixus*. This is the first report on the function of *Rp-BicC* in *R. prolixus* to provide a *bona fide* mechanism for egg formation.

In *D. melanogaster*, *BicC* is a maternal gene affecting embryonic anterior-posterior polarity, with a wide range of defects in segmentation [43,44]. We could not identify any embryonic structures in the eggs derived from *Rp-BicC* silenced females, as we have observed also for other maternal genes (Pagola, Pascual, and Rivera Pomar, unpublished data). This indicates that the role for *Rp-BicC* in embryogenesis, if any, has to be prior to gastrulation. Our results share similarities with the description of the *BicC* orthologue phenotype in the hemipteran *Nilaparvata lugens*, which seems to affect yolk loading in the egg [73]. Here we provide a plausible working hypothesis for the phenotype of *BicC* in the process, related to epithelial polarity as evidenced by the changes of distribution of endocytic pathway markers [74]. The physiology and biochemistry of yolk metabolism is well known in *R. prolixus* [41], however, the molecular and cellular mechanisms of egg formation and yolk accumulation are still scarce. It has been recently demonstrated that *Rp-ATG6* and *Rp-ATG8*, part of PI3P-kinase complexes that regulate the endocytic and autophagy machinery, are essential for yolk accumulation [71,72]. The phenotype of silenced *Rp-ATG6* females shares similarities with the one of *Rp-BicC*: unviable, small and white eggs that accumulate a minor fraction of yolk. However, *Rp-ATG6* seems to affect the oocyte uptake of yolk rather than the follicular cells, as they did not show defects in egg’s chorion. On the other hand, *Rp-ATG8* is required for the maternal biogenesis of autophagosomes and its role, although not exclusive, in follicular atresia. These results on the autophagocytic pathway and the ones presented here, point to common pathways affected by different genes at different levels.

Based on the expression of *Rp-BicC* in follicle cells and the distribution of vitellin and membrane markers in the silenced females, we support the idea that *Rp-BicC* affects the polarity of the follicular epithelium and, likely, the oocyte-follicle cell interaction. Cell-to-cell interactions are crucial for the development of oogenesis via proper yolk deposition and signaling from the follicle cell to the oocyte to establish embryonic polarity, the latter an aspect unknown in *R. prolixus* and worth to be further investigated. This suggests a conserved role for *BicC. Bicc1* (the mouse homologue of *BicC*) is required for E-cadherin-based cell-cell adhesion, indicating that that lack of *Bicc1* disrupts normal cell-cell junctions, and, in consequence, alter epithelial polarity [75].

Disruption of *BicC* in *D. melanogaster* affects the normal migration direction of the anterior follicle cell of the oocytes [46]. We observed that the primary effect of silencing *Rp-BicC* is a disorganized pattern of the follicular epithelium from the early previtellogenic stages until the end of vitellogenesis. At a first glance, the phenotype might be related to atresia. However, atresia, which can occur in any stage of oogenesis [35], results in a non-viable oocyte in which chorion deposition does not occur. The silenced *Rp-BicC* ovaries shows some characteristics of the atresia, such as the lack of a consistent perivitelline space between follicle cells, however, the follicle cells eventually produce the chorion. Although we cannot rule out that diminished egg production in the *Rp-BicC* silenced females is the consequence of increased atresia, the observation that all dissected ovaries showed the same atresic-like morphology in all oocytes, lead us to support the idea that atresia is not the main event. This differentiate the *Rp-BicC* phenotype from the *Rp-ATG6* and *Rp-ATG8* ones, although it requires further studies on the regulation of the different components of the pathways to shed light to the process. Interestingly, *BicC* has been described as a conserved translational regulator in animals and the available evidence indicates that it regulates many cellular processes [reviewed in 76].

New and exciting works on *R. prolixus* molecular and cellular mechanisms of oogenesis have open unexplored paths to understand the genetic and, therefore, the molecular interactions that regulate the formation of the egg. There is a challenge ahead for a more comprehensive understanding of the process of oogenesis in hemimetabolous insects. Deeper knowledge on this basic process in a vector of one of the most important disease in Latin America will pave the road to the design of new ways to control the population of the vector by affecting fertility.

## Acknowledgements

The authors thank all members of Rivera-Pomar and Andrés Lavore labs for fruitful discussions, L. Canavoso (CIBICI-CCT CONICET, Córdoba, Argentina) for kindly sharing the anti-vitellin antibody, A. Nazar for her contribution to develop the *in situ* protocol for *R. prolixus*, and Reprosemyx S.A. for kindly allowing the use of their confocal facility.

## Supporting information

**Fig S1. Multiple alignment of *BicC* orthologues.** Clustal W [77] was used to align sequences extracted from NCBI sequence database *Harpegnathos saltator* (gi|749730745-gi|749730739-X1), *Bombus terrestris* (gi|808147069), *Apis florea* (gi|820865347), *Megachile rotundata* (gi|805824678), *Acromyrmex echinatior* (gi|332022439), *Tribolium castaneum* (gi|91089717), *Pediculus humanus* (gi|242009357), *Drosophila melanogaster* (gi|24584541 and NP_723949.1), *Mus musculus* (AAK27347.1), *Nilaparvata lugens* [73], *Danio rerio* (NP_981965.1), *Xenopus laevis* (NP_001081996.1). The amino acid conservation is visualized with black blocks, the amino acid group level conservation with gray blocks. The dashes show the absence of sequence aligned along the alignment. Oblique lines refer to continuity of alignment.

**Table S1: Parental RNAi experimental data.**

